# FVIII interacts with cell surface to regulate endothelial cell functionality

**DOI:** 10.1101/2023.10.19.563105

**Authors:** Cristina Olgasi, Alessia Cucci, Ivan Molineris, Simone Assanelli, Francesca Anselmi, Chiara Borsotti, Chiara Sgromo, Andrea Lauria, Simone Merlin, Gillian Walker, Paola Capasso, Salvatore Oliviero, Antonia Follenzi

## Abstract

Haemophilia A (HA) is a rare bleeding disorder caused by factor 8 (F8) mutations. Clinical manifestations are spontaneous bleedings that primarily consist of hemarthrosis and intracranial haemorrhages. To date, the impairment of vessel stability in HA patients and the correlation between FVIII and endothelial functionality is poorly understood.

Here we show that FVIII plays a role in endothelial cell functionality. Blood Outgrowth endothelial cells (BOECs) knockout generated by CRISPR/Cas9, HA BOECs and HA iPSCs-derived ECs showed alteration of vessel-formation, endothelial cell migration, and vessel permeability. Importantly, the impaired EC phenotype was rescued by treatment with recombinant human FVIII or by lentiviral vector (LV) expressing FVIII. The FVIII function on endothelium was confirmed in vivo in a mouse model of severe HA which showed that an altered angiogenesis and vesselpermeability could be treated by exogenous FVIII. BOECstranscriptomic profiles revealed that FVIIIregulates the expression of endothelial basement membrane and extracellular matrix genes. Furthermore, exogenous expression of Nidogen2, identified as a FVIII regulated gene, restored the extracellular matrix integrity and EC functionality of HA ECs. In conclusion, FVIII is not only a coagulation factor but also an endothelial cell autocrine factor which promotes vessel stability.

## Introduction

FVIII gene (*F8*) is located on the X chromosome and when mutated, causes the X-linked disease of haemophilia A. The reduction or lack of plasma FVIII causes prolonged bleedings episodes, either spontaneously or secondary to trauma, with clinical severity proportional to the degree of FVIII reduction (1). To date, there is no definitive cure for HA, with the current therapeutic practice being a replacement therapy of recombinant human FVIII (rhFVIII) delivered by intravenous injections on an as-needs basis to treat acute haemorrhages or as prophylactic administration of FVIII to prevent bleeding events (2). Despite the amelioration with replacement therapy, 20 to 40% of patients with the severe form of HA develop inhibitors making the treatment ineffective (3, 4). Alongside clotting factor advancements, another class of drugs called non-replacement therapy is emerging to overcome the difficulties of intravenous delivery and to improve the effectiveness of therapies in all patients, regardless of the inhibitors formation (5–7). Bleeding events, however, can still occur after trauma requiring the use of additional haemostatic agents, according to the patients’ inhibitor status (8–10). Gene therapy strategies have been developed, with the ongoing clinical trials using adeno-associated viral vectors (AAVs), aimed at finding a definitive cure for HA (11–14).

Haemophilia A is associated with recurrent joint bleeding and intracranial haemorrhages (15, 16). Actually, one of the main and common long-term complications is the development of haemophilic arthropathy with joint impairment and chronic pain with a reduced quality of life starting from an early age (17–19).

Approximately 3 to 5% of severe HA patients develop intracranial haemorrhages (ICH) in the perinatal period, with trauma the major causative factor (20). The ICHs represent the most serious event that can occur in HA patients resulting in 20% rates of mortality and disability (21–23). The manifestation of ICH is often spontaneous, and it mainly occurs in patients that do not adhere to the prophylaxis regimen resulting in inadequate levels of blood FVIII (24, 25).

There are other important pathological conditions associated with FVIII deficiency, such as cartilage degeneration, bone remodelling and cardiovascular diseases (26–29). It has also been investigated the correlation between HA and cardiovascular disease development. In fact, despite the role of FVIII is not clear in the formation of atherosclerotic plaques, thrombin generation is likely to play a key role. Conversely, in terms of acute thrombosis, a risk factor for venous thromboembolism has been described in presence of supraphysiological FVIII levels (30).

The pathological mechanism underlying its development is not yet fully understood. The mechanism is multifactorial (31) and vascular development and angiogenesis are an essential component of blood-induced joint disease (32). Recently, it was demonstrated an attenuated microvascular endothelial functionality in haemophilic patients and altered collagen levels in the plasma of HA patients, suggesting a dysfunction in endothelial cells (ECs) (33, 34).

We and others have demonstrated that ECs are the main FVIII producers (35–37), however, an association between endothelial fragility and the absence of or low activity of FVIII has never been explored. Moreover, very little is known regarding the differences in the genetic profile between healthy and HA ECs. Healthy and HA ECs can be isolated from the peripheral blood, as blood outgrowth endothelial cells (BOECs) (38),or can be differentiated from induced pluripotent stem cells (iPSCs), reprogrammed from circulating CD34^+^cells (39). Both EC models, once transduced with a lentiviral vector (LV) carrying the B domain deleted (BDD) form of coagulation FVIII (LV- FVIII) and implanted into a prevascularized medical device, or in association with microcarrier beads,were able to correct the bleeding phenotype of HA mice (38, 39).

Therefore, based on the clinical observation of a general endothelial impairment in HA patients, here we aim to investigate the role of FVIII in the maintenance of ECs stability by evaluating the differences in the functionality and in the transcriptomic profile of healthy, HA and LV.FVIII- transduced HA ECs of both ECs models to define the involvement of FVIII in endothelial cell physiology and to investigate the potential extra-coagulative roles of FVIII in tissue homeostasis.

## Results

### ECs from haemophilic patients show impaired functionality

To investigate the role of FVIII in the maintenance of EC functionality, we evaluated the differences in the tubulogenesis of healthy (H), HA and LV-FVIII HA BOECs. H-BOECs formed a complete and stable network when cultured on Matrigel while HA-BOECs built a thin and incomplete network (Figure 1A). Importantly, treatment of HA-BOECs with rhFVIII, induced a significant improvement of the tubule network formation compared to non-treated HA-BOECs. Moreover, the FVIII secreted by LV-FVIII HA BOECs, showed a significant enhancement of vessel-like formation. Quantification of the formed network showed an increase in the number of nodes, junctions, branches, and total length of the tubules in HA-BOECstreated with rhFVIII or corrected by LV-FVIII (Figure S1A). To confirm FVIII involvement in EC functionality and stability, we also performed CRISPR/Cas9 knockout of *F8* in H-BOECs (KO-*F8* and Figure S1B-G) which showed a significantly decreased capability in forming tubule networks in Matrigel (Figure 1B and S1H). Importantly, the rescue of FVIII by treating KO-*F8* cells with rhFVIII, induced an increase in the tubule network.

**Figure 1.**
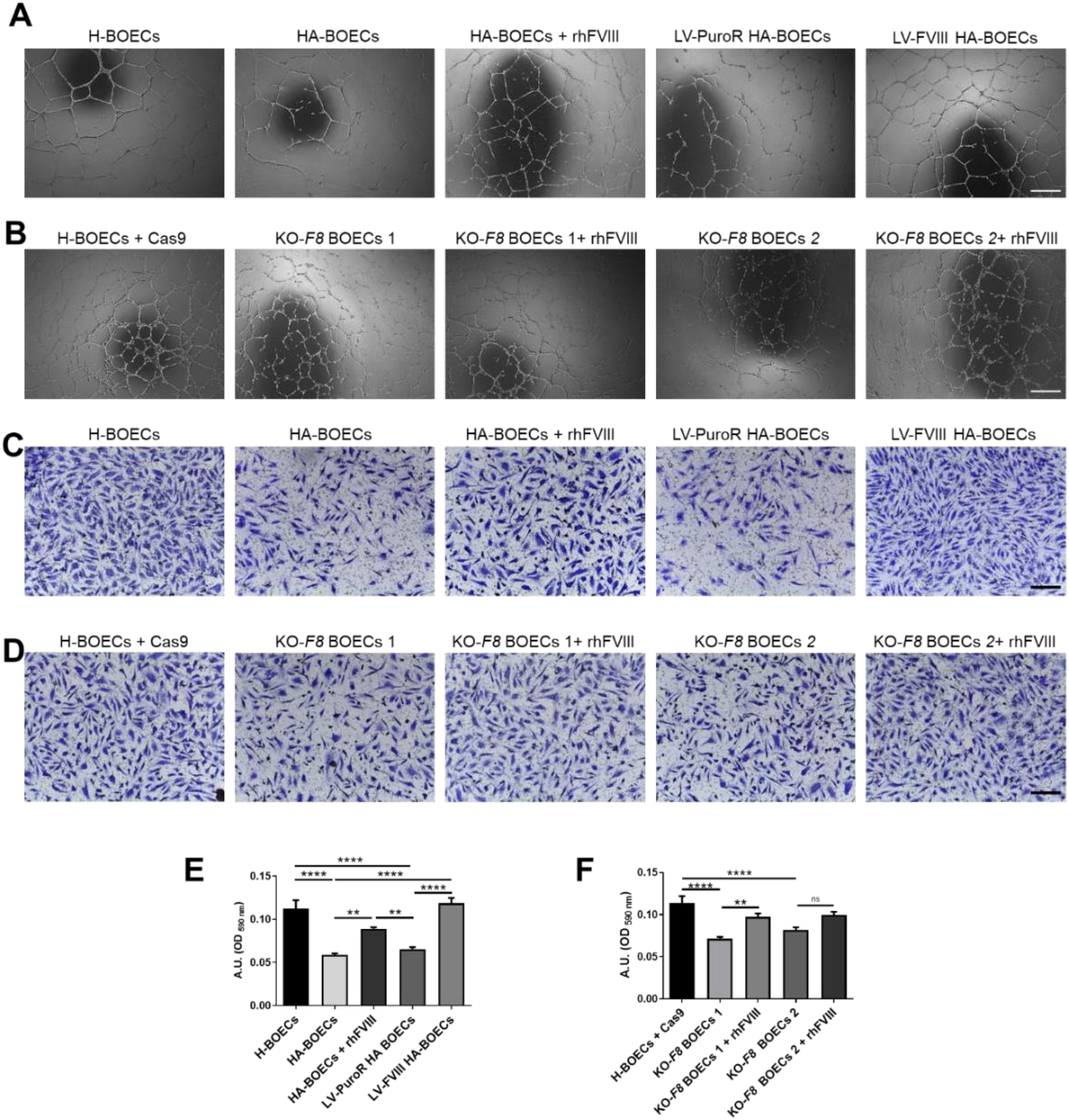
HA-BOECs are defective in tubulogenesis and migration. **A)** Representative images of tubulogenic assay on healthy (n=4), HA (n=4) and in presence or absence of rhFVIII, and LV-FVIII HA-BOECs (n=4). LV-PuroR was used as control for LV transduction. Scale bar = 500 µm **B)** Representative images of tubulogenic assay on H-BOECs LV transduced with Cas9 (H-BOECs + Cas9), KO-*F8* BOECs 1 and 2, in presence or absence of rhFVIII. Scale bar = 500 µm **C**) Representative images of migrated cells on H-BOECs, HA-BOECs, HA-BOECstreated with rhFVIII and LV-FVIII HA-BOECs. LV-PuroR was used as control for LV transduction. Scale bar = 200 µm. **D)** Representative images of migrated cells on H-BOECs+ Cas9, KO-*F8* BOECs 1 and 2, in presence or absence of rhFVIII. Scale bar = 200 µm. **E)** Indirect quantification of cell migration assay by elution of crystal violet staining on H-BOECs, HA-BOECs, HA-BOECs treated with rhFVIII and LV-FVIII HA-BOECs. LV-PuroR was used as control for LV transduction. **F)** Indirect quantification of cell migration assay by elution of crystal violet staining on H-BOECs + Cas9, KO-*F8* BOECs 1 and 2, in presence or absence of rhFVIII. (****p < 0.0001***p < 0.001; ** p 0.01; * p < 0.05). Data are expressed as mean ± SD and are representative of three independent experiments.

Next, we analysed the EC capability to migrate. HA-BOECs and KO-*F8* BOECs showed a reduced migration capability which was significantly recovered by treatment with rhFVIII and by LV-FVIII transduction (Figure 1C-F).

The ability to recover their angiogenic capacity and migration was also observed in iPSC-derived ECs obtained from HA patients (HA-iECs) (39) and in KO-*F8* HMEC-1 (Figure S1B-G) treated with rhFVIII or by LV-FVIII compared to non-transduced HA-iECs (Figure S2 and S3).

Taken together, the above data demonstrate a novel function for FVIII which, in addition to its known role in coagulation, plays a direct role in promoting endothelial cell functionality.

### FVIII drives *in vivo* vessel formation and stabilization

Based on the *in vitro* results we further investigated FVIII angiogenic activity *in vivo*. We first assessed the angiogenic potential in NOD Scid IL2R^γnull^ (NSG) mice versus NSG-HA mice by intradermal implantation of Matrigel plugs that were removed after 10 days and analysed by immunofluorescence. The NSG mice showed well organized CD31 positive vessels surrounded by pericytes stained with αSMA (Figure 2A). In contrast, NSG-HA mice formed small and disorganized vessels (Figure 2B). We then intradermal injected NSG-HA mice with Matrigel containing rhFVIII. Remarkably, vessels formed within Matrigel plugs supplemented with rhFVIII showed a well-organized structure (Figure 2C) compared to Matrigel plugs without rhFVIII. The effect of FVIII on neo angiogenesis in Matrigel plugs was also evident in mice treated by gene therapy (5×10^8^ TU/mouse of LV-FVIII), which showed the formation of larger and stable vessels (Figure 2D). Quantification of vessel density and vessel diameter confirmed the ability of FVIII to promote *in vivo* angiogenesis (Figure 2E-F). These results demonstrate that FVIII plays an *in vivo* functional role in vessel formation.

**Figure 2.**
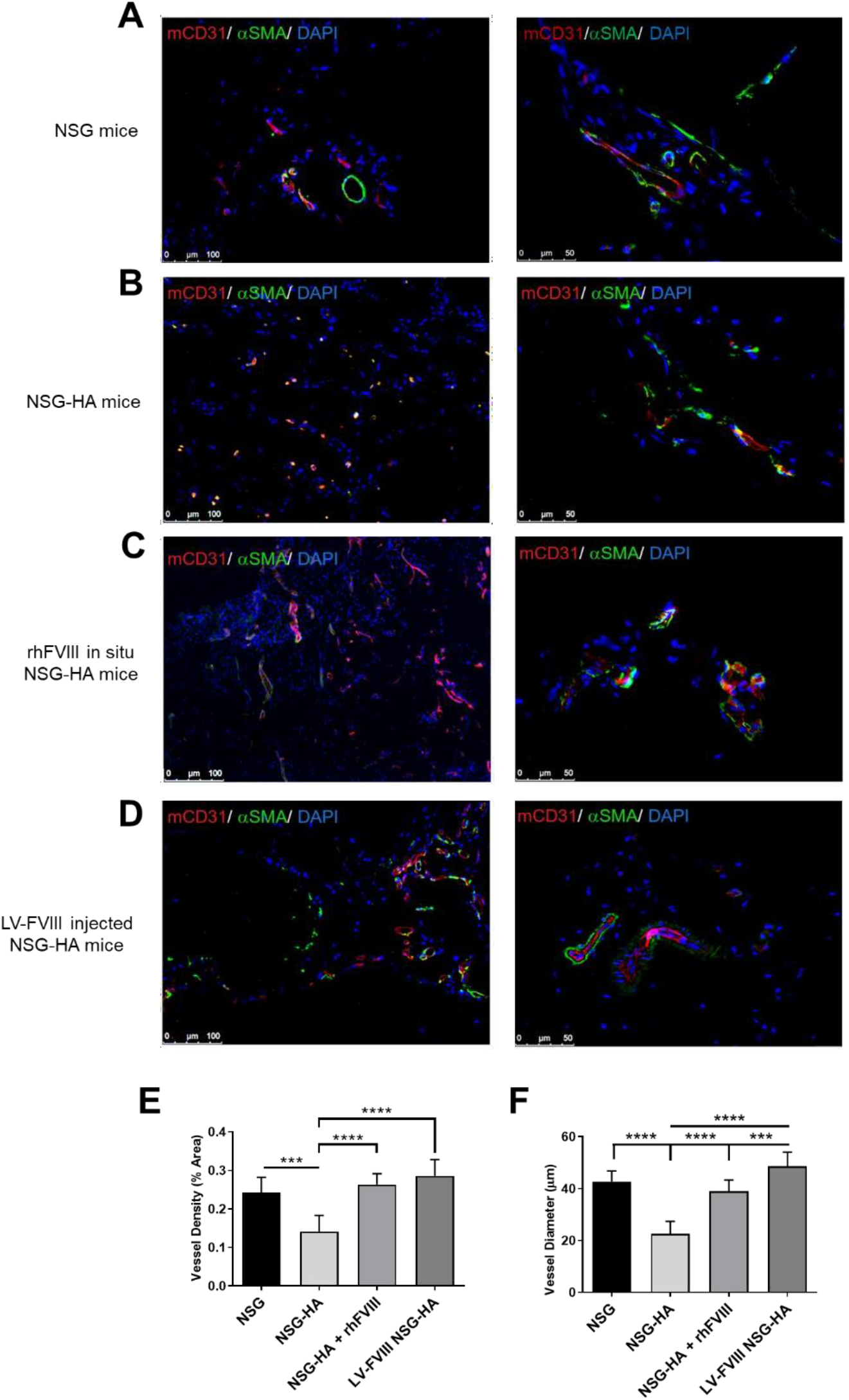
FVIII is required for vessel migration *in vivo* in HA mice. Immunofluorescence staining for mCD31 (red) and αSMA (green) on vessels formed within Matrigel plug harvested from: **A)** NSG mice, **B)** NSG-HA mice, **C)** NSG-HA mice treated *in situ* with rhFVIII, **D)** NSG-HA mice injected with LV- FVIII. Data are representative of three independent experiments (Total number = 9 for each condition). **E)** Quantification of vessel density in Matrigel plugs. **F)** Quantification of vessel diameter in Matrigel plugs. (****p < 0.0001***p < 0.001). Data are expressed as mean ± SD.

### FVIII drives *in vivo* vessel formation and stabilization of injected human ECs

We next analysed whether human BOECs, expressing GFP (Figure S4A) embedded in Matrigel plugs could also form vessels *in vivo* in NSG-HA mice. Indeed, 10 days after their implantation H-BOECs showed a well-organized vessel morphology in Matrigel plugs (Figure 3A) while HA-BOECs failed to form vessels (Figure 3B). Importantly, the administration of rhFVIII into the Matrigel plugs significantly improved the ability of HA-BOECs to form well-organized vessels (Figure 3C). Likewise, LV-FVIII HA-BOECsdisplayed improved vessel formation (Figure 3D) compared to HA- BOECs. Vessel density and diameter quantification confirmed that rhFVIII and LV-FVIII transduction significantly ameliorated the ability of human ECs to form well organized vessels in Matrigel plugs implanted in HA mice (Figure 3E-F).

**Figure 3.**
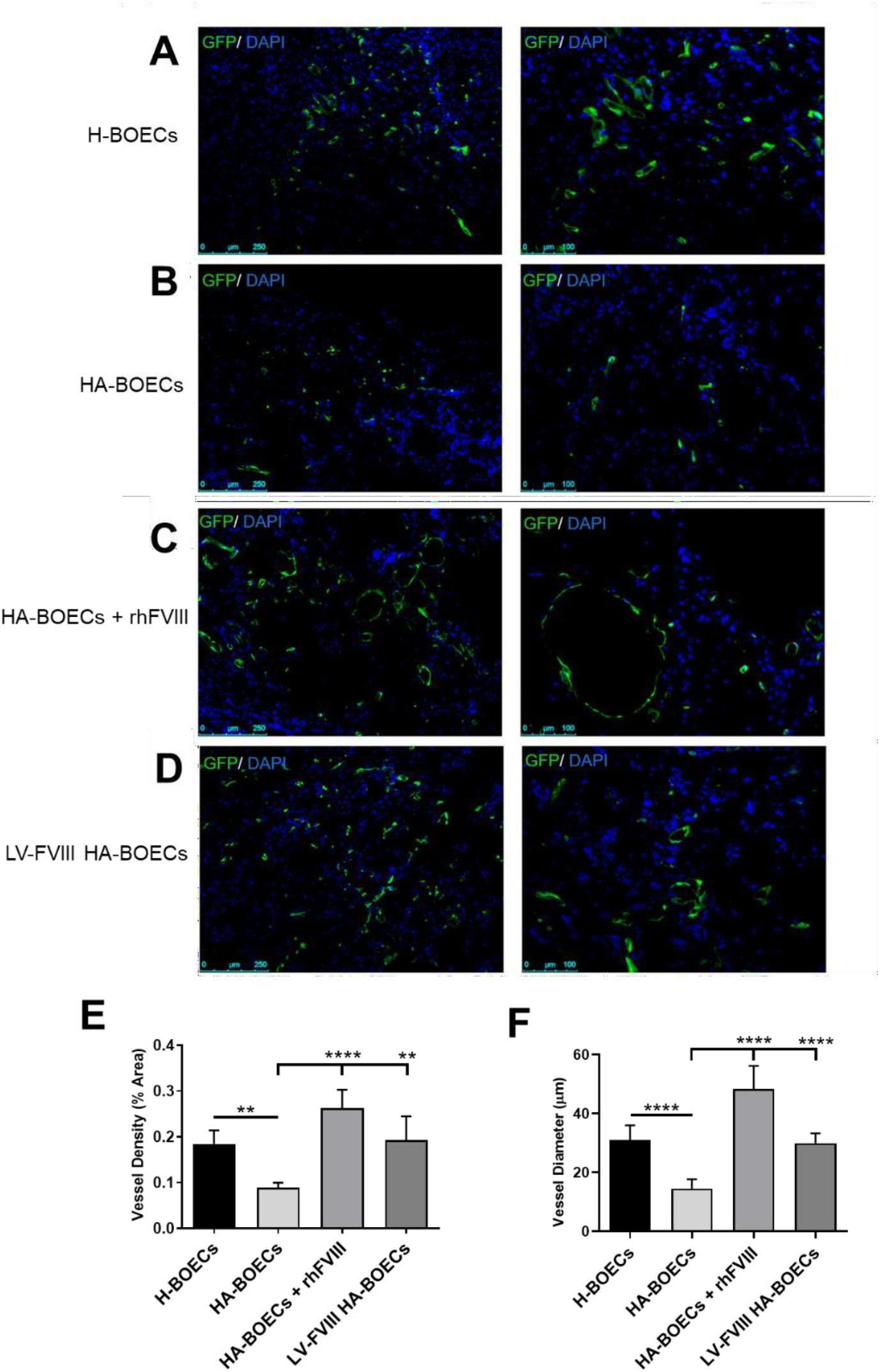
FVIII enhances angiogenic potential of HA-BOECs *in vivo.* Immunofluorescence detection of GFP+: **A)** H-BOECs, **B)** HA-BOECs, **C)** HA-BOECs treated with rh-FVIII, **D)** LV- FVIII HA-BOECs, forming vessels within Matrigel plugs transplanted intradermally in NSG-HA mice. (Total number = 9 each condition). **E)** Quantification of vessel density. **F)** Quantification of vessel diameter (****p < 0.0001; ** p 0.01). Data are expressed as mean ± SD.

### Lack of FVIII expression in ECs alters vessel permeability both *in vitro* and *in vivo*

As the endothelium serves as a barrier between circulating blood and the surrounding tissues, we also analysed the role of FVIII in vessel permeability by measuring the extravasation of FITC-conjugated dextran through an EC monolayer plated in Transwells. Quantification of permeate fluorescence showed an enhanced permeability of HA-BOECs compared to healthy cells (Figure 4A). The same pattern was observed in KO-*F8* BOECs (Figure 4B). Furthermore, both HA and KO-*F8* ECs treated with rhFVIII or corrected by LV-FVIII significantly reduced the dextran leakage (Figure 4A, B). These results were also confirmed in HA-iECs and KO-*F8* HMEC-1 (Figure S2E, S3E respectively), therefore indicating that FVIII is required to inhibit EC permeability.

**Figure 4.**
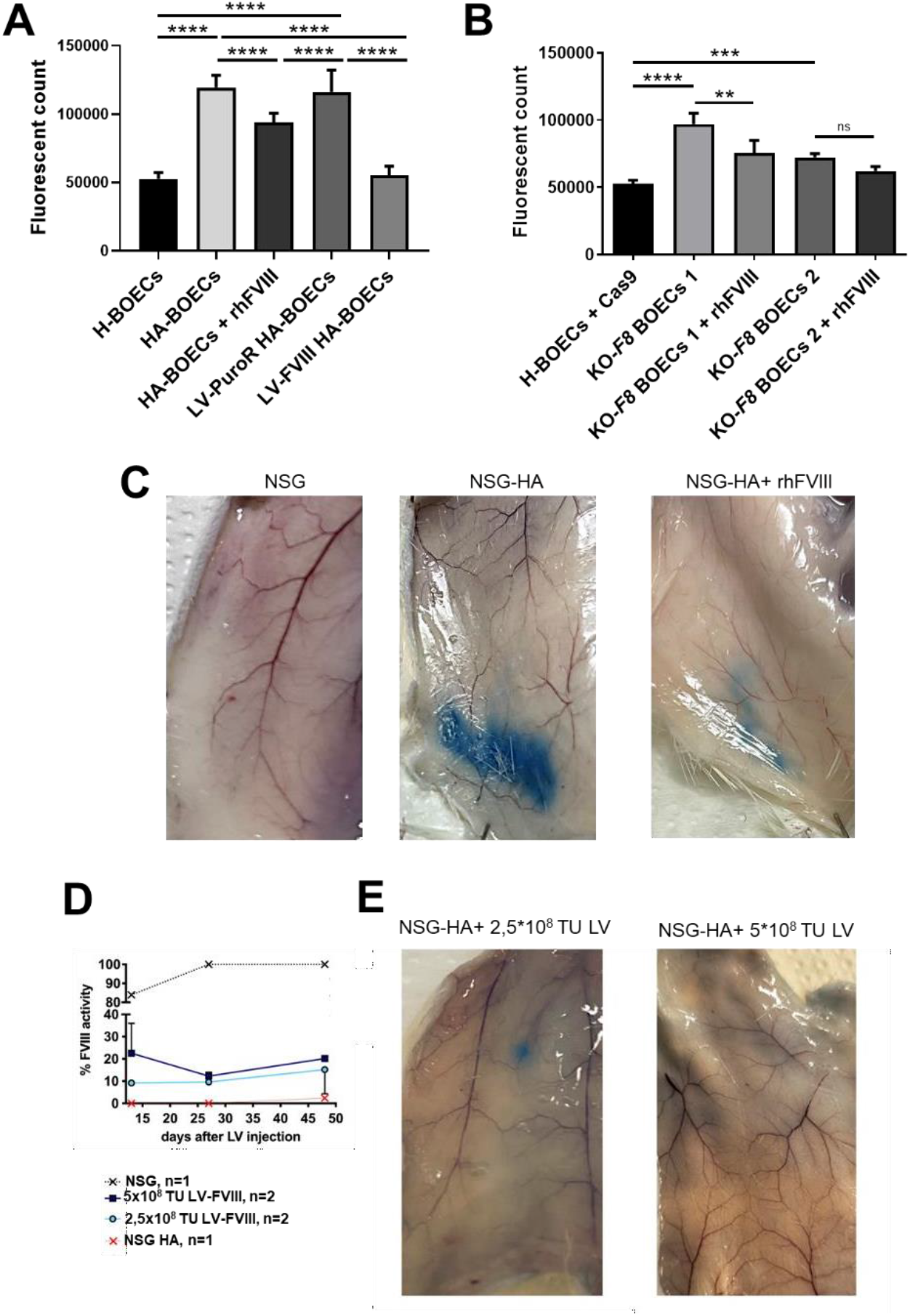
FVIII regulates vascular permeability. **A)** *In vitro* permeability assay quantification calculated on the extravasation of FITC-dextran through an intact monolayer of H-BOECs, HA- BOECs, HA-BOECstreated with rhFVIII and LV-FVIII HA-BOECs. LV-PuroR was used as control for LV transduction. **B)** Quantification of the extravasation of FITC-dextran on H-BOECs + Cas9, KO-*F8* BOECs 1 and 2, in presence or absence of rhFVIII. **C)** Representatives picture of Evans Blue dye extravasation in interstitial tissues of NSG, NSG-HA mice, NSG-HA treated with rhFVIII. **D)** Kinetics of the percentage of FVIII activity measured by aPTT assay in the plasma of NSG-HA mice tail vein injected with 2,5×10^8^ or 5×10^8^ TU of LV-FVIII. **E)** Macroscopically evaluation of Evans Blue dye distribution under the skin in NSG-HA injected mice with 2,5×10^8^ or 5×10^8^ TU of LV- FVIII.

To analyse *in vivo* vessel permeability, NSG and NSG-HA mice were intravenously injected with the Evans Blue albumin-binding dye. Under physiologic conditions the endothelium is impermeable to albumin and therefore Evans Blue remains restricted within blood vessels(40) (Figure 4C, left panel). In contrast, in NSG-HA mice, the endothelium becomes permeable to small proteins such as albumin allowing the extravasation of Evans Blue in the surrounding tissues (Figure 4C, central panel). Importantly, mice treated every 2 days with 4 IU of rhFVIII for 20 days, restored physiological vascular permeability as shown by the significant reduction of Evans Blue extravasation (Figure 4C, right panel).

This effect was also evident after tail vein injection in NSG-HA mice of two different dosages (2.5×10^8^ TU or 5×10^8^ TU/ mouse) of LV-FVIII. The FVIII activity was evaluated over time for up to 50 days showing that FVIII in injected mice induced a marked reduction of dye extravasation (Figure 4D, E), demonstrating that FVIII is required for *in vivo* to control vessel permeability.

### Transcriptomic profile identified genes regulated by FVIII in ECs

To study the impact of FVIII on the endothelial cell transcriptome, we performed RNA-seq on H- BOECs, HA-BOECs, and LV-FVIII HA-BOECs.

Volcano plot identified approximately 150 genes downmodulated in HA versus (vs) H BOECs (Figure 5A) of which 51 were significantly rescued after LV-FVIII transduction (Figure 5B, C). Among the rescued genes we identified EC response to signalling such as AKAP12 (41) and the deacetylase HDAC9 (42) as well as endothelial surface proteins including integrin ITGAV (43) and the basement membrane components Nidogen 2 (NID2) (44), collagen IVa1 (45) and PXDN (46).

**Figure 5.**
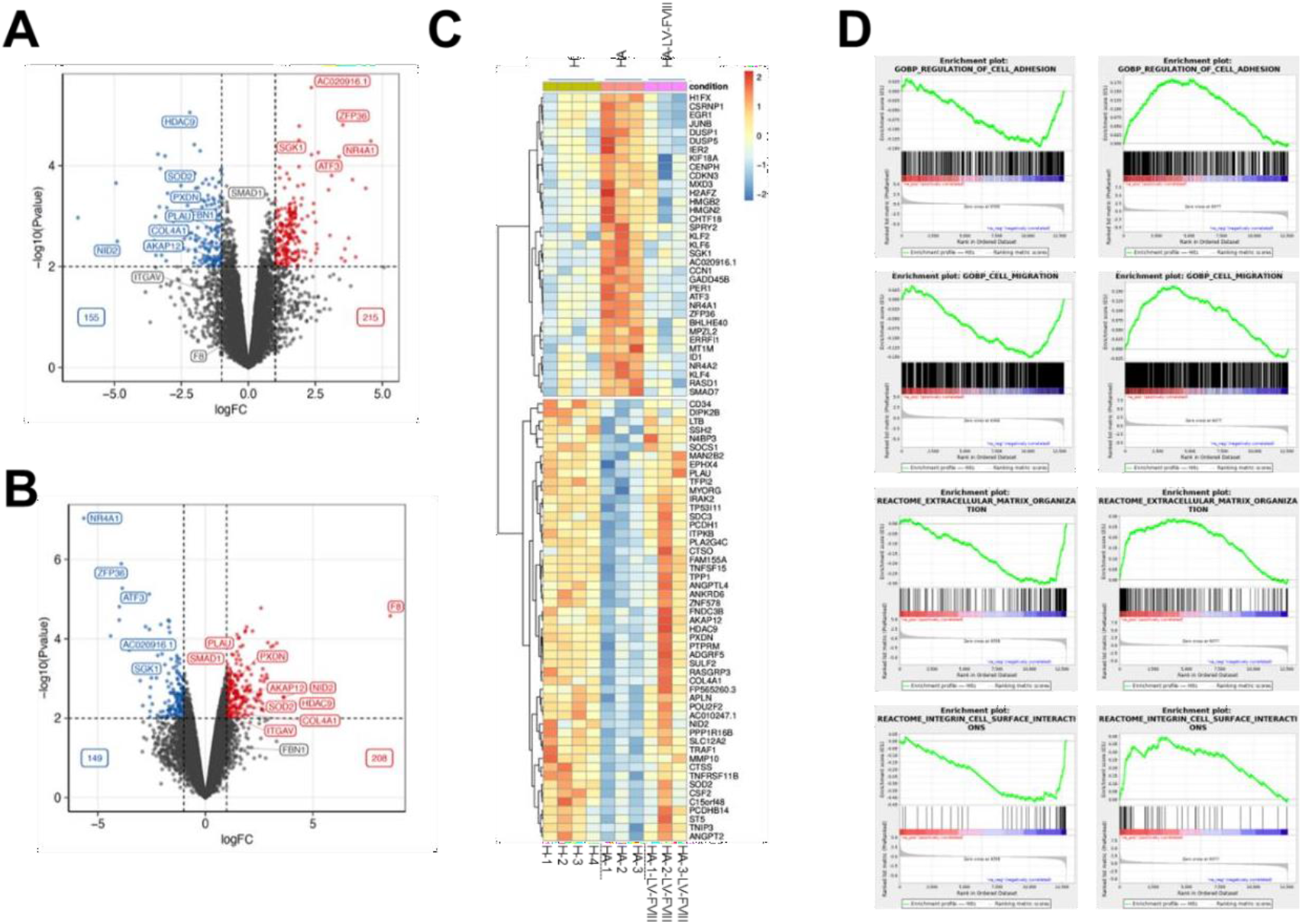
Genes regulated by FVIII. **A)** Volcano plot showing differentially expressed genes in HA vs H-BOEC (disease). Up-regulated: log fold change > 1 and p value <0.01. Down-regulated: log fold change < -1 and p value <0.01. **B)** Same as A but for transduced BOEC vs HA (transduction). **C)** Heatmap showing the expression pattern of genes rescued by the transduction, i.e. genes that are significantly differentially expressed both in disease and in transduction, but change the sign of logFC: down-regulated in the disease and up-regulated in the treatment or up-regulated in the disease and down-regulated in the transduction. **D)** GSEA plots of pathways that resulted downregulated in disease and rescued by the transduction.

In agreement, Gene Set Enrichment Analysis (GSEA) of the differentially expressed genes, identified pathways corresponding to cell adhesion, cell migration, extracellular matrix organization, and integrin cell surface interaction which were down-modulated by the disease and rescued by LV-FVIII transduction (Figure 5D).

### NID2 rescues the FVIII EC functionality

Our data suggest that FVIII signalling regulates genes required for the endothelial basement membrane integrity. To gain insight into the functional defects of ECs in the absence of a FVIII, we focused on NID2, a glycoprotein involved in the endothelial basal membrane stability, which was significantly down regulated in haemophilic cells and rescued by LV-FVIII transduction (Figure 6A). We therefore performed a complementation assay through the ectopic expression of NID2 in HA BOECs (Figure S4B). The expression of NID2 in HA-BOECs induced the formation of a stable tubule network in Matrigel (Figure 6B, C). Migration and permeability assays showed that NID2 ectopic expression in HA-BOECs rescued the migration capability and reduced the permeability (Figure 6D- F) to a level like H-BOECs. To further confirm the interplay between FVIII and NID2 weperformed a NID2 knockdown by specific shRNA in H-BOECs and HMEC-1 (Figure S4C, S4D).We observed that NID2 down-modulation significantly reduced EC tubulogenesis (Figure 7A, B andS5A, S5B), migration (Figure 7C, D and S5C, S5D) and increased permeability (Figure 7E and S5E).Interestingly, the treatment of shNID2 H-BOECs and HMEC-1 with rhFVIII did not improve any of the EC functions (Figure 7 and S5) confirming that NID2 is a downstream effector of FVIII.

**Figure 6.**
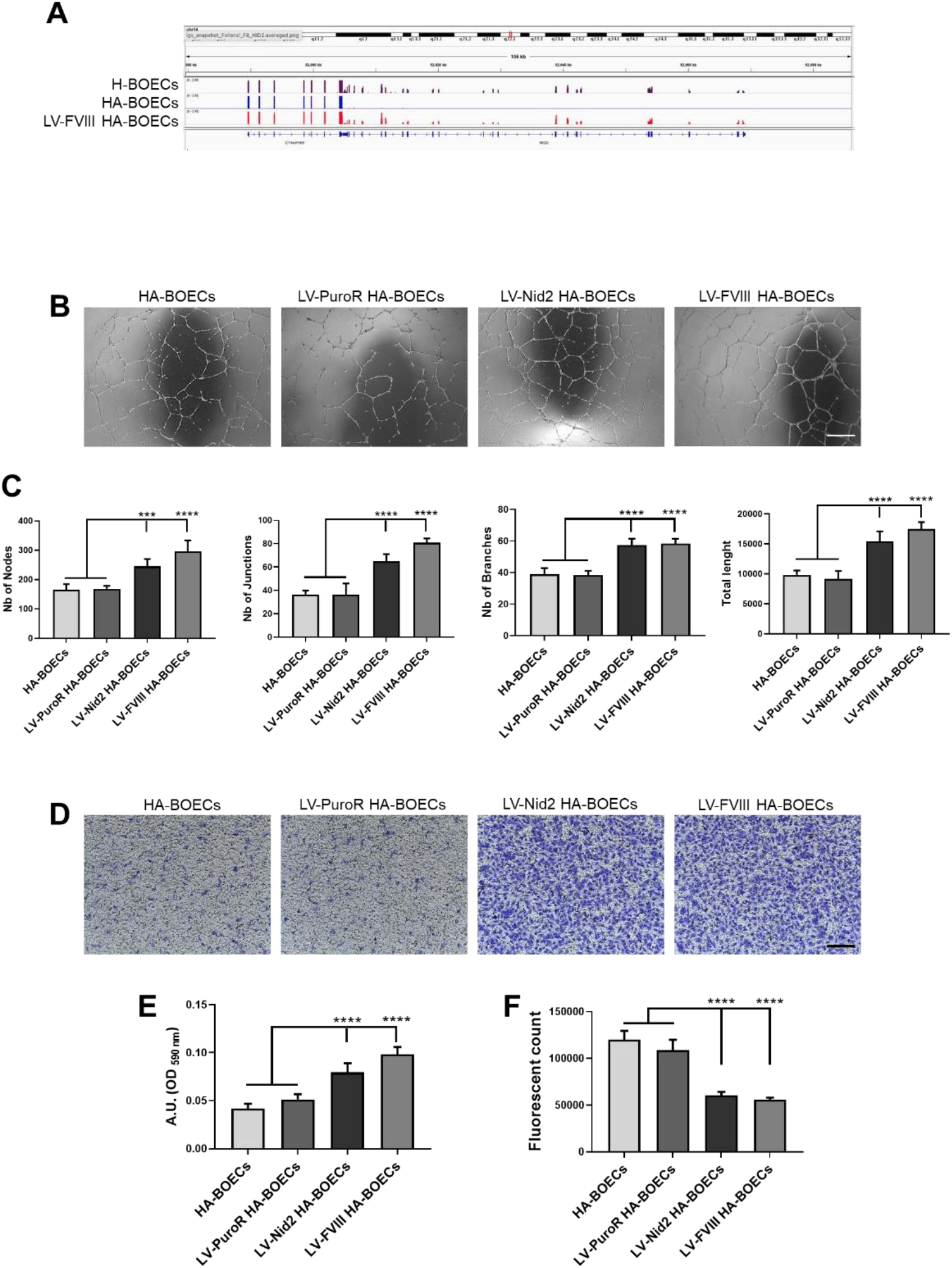
NID2 rescues the FVIII deficiency in EC functionality. **A)** Genome browser view showing the RNA-seq signal profile of the NID2 gene in H-BOECs, HA- BOECsand LV-FVIII HA-BOECs. **B**) Representative images of tubulogenic assay on HA, LV-NID2 and LV-FVIII HA-BOECs. LV-PuroR was used as control for LV transduction. Scale bar = 500 µm. **C)** Quantification of number of nodes, junctions, branches and total length of the tubule networks. **D**) Representative images of migrated cells on HA, LV-NID2 and LV-FVIII HA-BOECs. LV-PuroR was used as control for LV transduction. Scale bar = 200 µm. **E)** Indirect quantification of cell migration assay by elution of crystal violet staining. **F)** Permeability assay quantification calculated on the extravasation of FITC-dextran through an intact monolayer of HA, LV-NID2 and LV-FVIII HA-BOECs. LV-PuroR was used as control for LV transduction. (****p < 0.0001***p < 0.001). Data are expressed as mean ± SD and are representative of three independent experiments.

**Figure 7.**
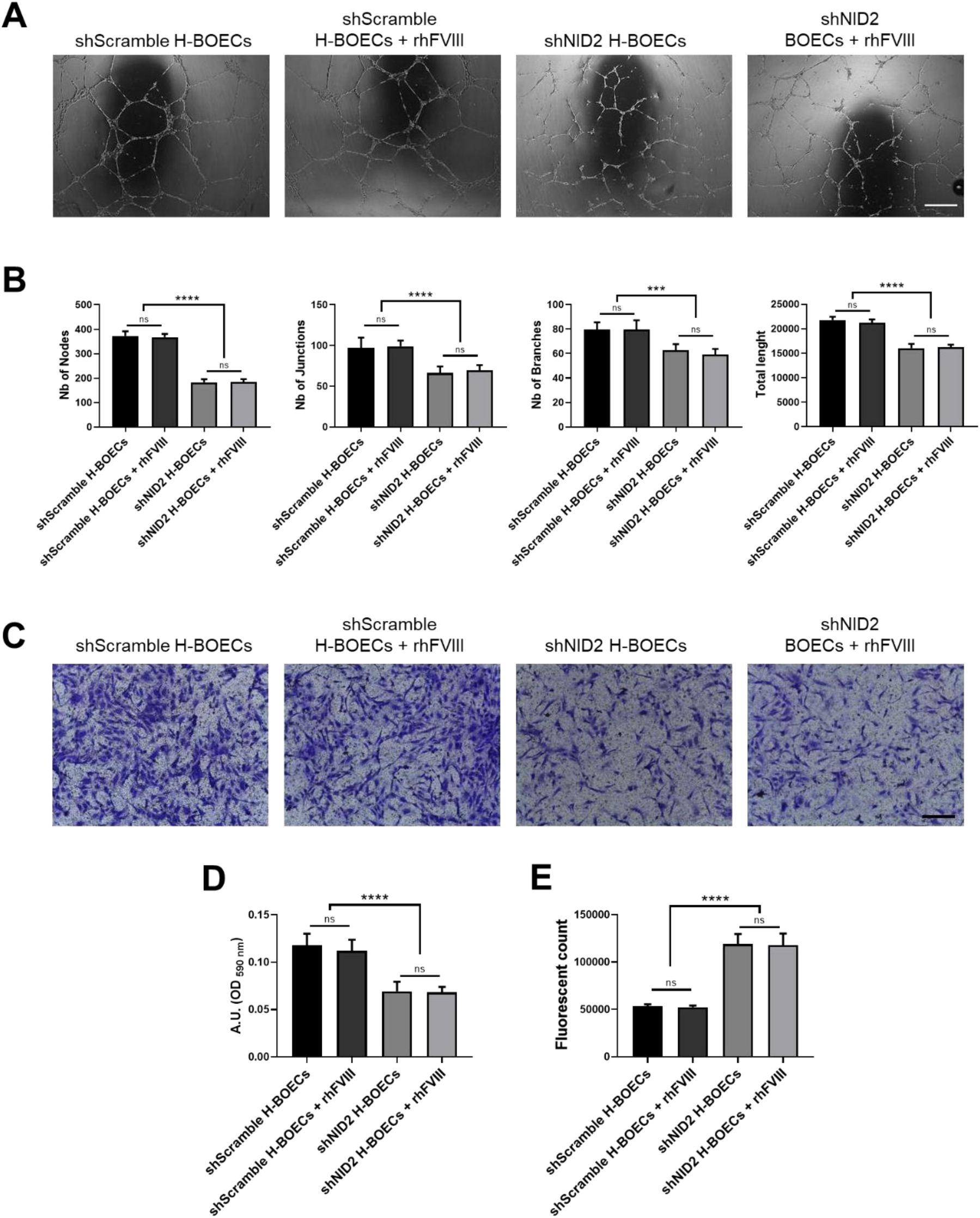
NID2 knockdown impairs H-BOECs endothelial functionality. **A**) Representative images of tubulogenic assay on shScramble and shNID2 H-BOECs in presence or absence of rhFVIII. Scale bar = 500 µm. **B**) Quantification of number of nodes, junctions, branches and total length of the tubule networks. **C**) Representative images of migrated cells on shScramble and shNID2 BOECsin presence or absence of rhFVIII. Scale bar = 200 µm. **D**) Indirect quantification of migrated cells by elution of crystal violet staining. **E**) Permeability assay quantification calculated on the extravasation of FITC-dextran through an intact monolayer of shScramble and shNID2 BOECs in presence or absence of rhFVIII. (****p < 0.0001 ***p < 0.001) Data are expressed as mean ± SD and are representative of three independent experiments.

## Discussion

FVIII is well recognized as an important plasma factor involved in blood clotting as its mutations lead to HA, a recessive X-linked bleeding disorder caused by coagulation defects (47). Here we have demonstrated that, in addition to its clotting function, FVIII also plays a regulatory role in endothelial cell functionality. Increased vascular permeability is one of the major indicators of blood vessel damage in haemophilic patients. It has been reported to be associated with hemarthrosis that could lead to recurrent joint bleedings (48, 49), in addition to being a risk factor for spontaneous brain bleedings(21).

By several angiogenic assays we show that endothelial cells from HA patients and KO-*F8* ECs, failed *in vitro* and *in vivo* angiogenic tests demonstrating the extra-coagulative role of FVIII. The presence of rhFVIII embedded in the Matrigel plug *in vivo* in NSG-HA mice significantly improved the ability of haemophilic mouse ECs to migrate in the Matrigel and form well-organized vessels with structured lumen formations, suggesting that FVIII is required for angiogenesis. The endothelial dysfunction was restored by in situ delivery of rhFVIII or systemic LV-FVIII gene therapy inducing a significant increase in vessel density and diameter compared to NSG-HA mice injected with Matrigel alone. The constant presence of exogenous human FVIII in the matrix led to the formation of newly organized vessels.

Interestingly, the presence of rhFVIII embedded in the Matrigel plug in NSG-HA mice greatly also improved the ability of HA BOECs to form well-organized vessels with structured lumen formation, confirming that FVIII is required for neoangiogenesis of human endothelial cells.

Moreover, lack of FVIII in vitro and in vivo increased vessel permeability with the EC functionality rescued both by rhFVIII and LV-FVIII delivery, demonstrating that FVIII is required to recover the physiological vessel permeability. Importantly, 10 to 20 % of FVIII activity in plasma of LV-FVIII corrected haemophilic mice was sufficient to restore vascular permeability. Increased vascular permeability is one of the major indicators of blood vessel damage in hemophilic patients(48, 49). It has been reported to be associated with hemarthrosis that could lead to recurrent joint bleedings, in addition to being a risk factor for spontaneous brain bleedings(21). Therefore, our results support the hypothesis of a general vascular fragility in HA patients that can lead not only into joint bleedings but also to the uncanonical clinical manifestations. Thus, FVIII is an endothelial factor whose activity is required not only for blood coagulation but also to maintain endothelial homeostasis. To our knowledge, the correlation between endothelial fragility and the absence or low activity of FVIII has never been explored and very little is known about the differences in the genetic profile between healthy and hemophilic endothelial cells.

To gain insight into the endothelial effectors of FVIII in the maintenance of EC stability, we analysed the transcriptome of healthy, KO-*F8*, and rescued BOECs. The expression patterns highlighted significant changes between healthy and HA ECs which were rescued by LV-FVIII correction. GSEA of the differentially expressed genes, identified pathways corresponding to cell adhesion, cell migration, and integrin cell surface interaction with extracellular matrix that are down-modulated in HA BOECs and rescued by LV-FVIII transduction.

This transcriptomic analysis can therefore explain the differences in endothelial functionality, as migration, tubulogenesis and permeability observed both *in vitro* and *in vivo*. Interestingly, the impairment observed in our *in vitro* and *in vivo* experiments in HA EC functionality can be also correlated to the down-modulation of genes related to ECM, which play a fundamental role in the stabilization of ECs. ECM has key roles in tissue remodeling, in the maintenance of tissue integrity and importantly, is involved in the maintenance of the organization of vascular ECs into blood vessels(50). Migration capability of ECs is related to their adhesion to ECM and is essential for angiogenesis and sprouting of blood vessels(51).

Among the differentially expressed genes identified by RNAseq, we focused on NID2 a gene particularly interesting as it codes for a cell adhesion glycoprotein required for endothelial basement membrane integrity stabilizing collagen IV and laminin network (44, 52). Studies investigating nidogen’s role in modulating cellular phenotypes and differentiation are still in the early stages and knowledge about how nidogens regulate intracellular pathways remains limited (44, 53, 54). The reduction of NID2 in the basal membrane of endothelial cells in hemophilic patients could explain an increased vessels fragility, which, in stress conditions, might result in blood leakage. Our experiments revealed that NID2 is a downstream effector of FVIII in endothelial functionality as we found that NID2 was significantly downregulated in HA-ECs and rescued by FVIII while FVIII treatment did not rescue *NID2* knockdown confirming that NID2 is downstream of FVIII in endothelial cells. Importantly, *NID2* knockdown resulted in ECs impairment similarly to what we observed in HA ECs and its ectopic expression rescued tubulogenesis, migration, and permeability of haemophilic ECs.

Taken together our data demonstrate that lack of FVIII in HA patients impairs not only coagulation but also endothelial transcriptome predisposing patients to increased risk of trauma due to vessel fragility, confirming that FVIII is an endothelial cell autocrine factor which promotes vessel stability.

## Materials and Methods

### Cell culture

iPSC-derived ECs (iECs) were differentiated as previously described (39) from iPSCs reprogrammed from CD34^+^ cells isolated from peripheral blood of two healthy donors and two severe HA patients. BOECs were isolated from four healthy donors and four severe HA patients as previously described(38).

### *In vitro* tubulogenic assay

Twenty-four-well tissue culture plates were coated with 300 µl Matrigel Matrix (Corning) per well and allowed to solidify at 37°C for 30 minutes. 5×10^4^ BOECs or 5×10^4^ HMEC-1 or 3×10^5^ i-ECs, were resuspended in culture medium and placed on top of the Matrigel. Plates were incubated at 37°C, 5% CO2 and analysed after overnight (ON) incubation. For treatment with recombinant human BDD FVIII (rhFVIII) cells were incubated on top of Matrigel ON with 1U/ml of rhFVIII. Images were acquired under inverted microscope Leica ICC50. ImageJ Angiogenesis software was used for the quantification of number of nodes, junctions, branches, and total length.

### *In vitro* migration assay

BOECs or HMEC-1 or iECs were plated into the upper compartment of 8-μm pore size Transwell (Corning) at a density of 10^5^ cells in serum-free medium while the lower compartment of the chamber was filled with complete medium supplemented with VEGF-A (50 ng/ml) and incubated ON. FVIII- treated cells were incubated on top of transwell ON with 1U/ml of rhFVIII. Afterincubation, migrated cells were fixed with 70% ethanol and stained with 0,1% Crystal violet (Sigma-Aldrich). The migrated cells were photographed under inverted microscope Leica ICC50. Crystal violet was eluted with 10% acetic acid and quantified using Victor Spectrophotometer (Hi-Tech Detection Systems) at 590 nm.

### *In vitro* permeability assay

Permeability was measured across a monolayer of BOECs or HMEC-1 or iECs. Cells (80,000 cells/well) were plated on 0.1% gelatin coated Transwell (8 μm pore, 24-well format, Corning) and cultured until confluence was reached. FVIII-treated cells were incubated on top of transwell with 1U/ml of rhFVIII added every 2 days. At the end of the culture, 50 μl (5 μg/ml) of FITC-conjugated 40-kDa dextran (Sigma-Aldrich) was added to the upper chamber and the fluorescence of the lower chamber was measured in the medium after 30 min of incubation using Victor Spectrophotometer at 490 nm (excitation)/520 nm (emission). Fluorescence readings were normalized to dextran permeability in transwell inserts without cells.

### RNA-seq

RNA from the different ECs was purified as previously described (55) and its integrity was measured using Fragment Analyzer™ (Advanced Analytical). Library preparation was performed from PolyaAplus RNA using Illumina TruSeq RNA prep-kit as previously described. Samples were run in the Illumina sequencer NextSeq 500.

Sequencing reads were aligned to human reference genome (version GRCh38.p13) using STAR v2.7.7a0 (56) (with parameters –outFilterMismatchNmax 999 –outFilterMismatchNoverLmax 0.04) and providing a list of known splice sites extracted from GENCODE comprehensive annotation (version 32) (57). Gene expression levels were quantified with featureCounts v1.6.3 (58) (options: -t exon -g gene_name) using GENCODE gene annotation (version 32 basic). Multi-mapped reads were excluded from quantification. Gene expression counts were next analyzed using the edgeR package (59). Normalization factors were calculated using the trimmed-mean of M-values (TMM) method (implemented in the calcNormFactors function) and RPKM were obtained using normalized library sizes and gene lengths. After filtering the lower expressed genes, a differential expression analysis was carried out by fitting a GLM to all groups and performing LF test for the interesting pairwise contrasts, blocking on patients when possible. Genes were considered as significantly differentially expressed (DEGs) when having |logFC| >1 and raw p value < 0.01 in each reported comparison, as advised by SEQC consortium (60).

After the identification of a dataset of differentially expressed genes (DEGs), Enrichr online tool was used to identify pathways and gene ontology (GO) terms enriched using DEGs as input. A term is defined as significantly enriched if the reported adjusted p value is < 0.01.

Gene Set Enrichment Analysis (GSEA) was performed using Broad Institute java package version 3.0 (classic mode) and MSigDB version 7.5.1 (61).

### CRISPR/Cas9 two vector system

Healthy BOECs and HMEC-1 cells were LV transduced with a two vectors CRISPR/Cas9 system(62, 63). The first vector contains S. pyogenes Cas9 nucleotide sequence under the control of a doxycycline synthetic promoter and the Puromycin resistance gene under the control of human PGK promoter (Figure S1B). Transduced cells were selected with a puromycin (1ug/ml; Sigma) enriched medium prior the second LV transduction. The second vector carries the gRNA sequence against F8 gene under the control of U6 promoter and the GFP under the control of human PGK promoter was used as a second selection marker (Figure S1B). Two gRNAs against the F8 gene were chosen amongst the six designed (Figure S1C). Cas9 expression was induced with Doxycycline (Sigma) at 75ng/ml for 72 hours.

### In vivo permeability assay

Evans Blue extravasation was used to quantify the capillary permeability in 8 week old NSG (NOD.Cg-*Prkdc^scid^ Il2rg^tm1Wjl^*/SzJ, Jackson #005557) and NSG-HA mice (Italian Health Ministry Authorization nos. 492/2016-PR and DBO64) (37). For FVIII delivery 4 IU/mice of rhFVIII were tail vein injected every 2 days for 20 days. NSG-HA mice were also tail vein injected with two different dosages (2.5×10^8^ TU or 5×10^8^ TU/ mouse) of LV-FVIII. The FVIII activity was evaluated over time for up to 50 days. At the end of the experiment, a 0.5% Evans Blue solution (Sigma-Aldrich) was injected into the tail vein. After 15 minutes, the mice were killed, and the extravasation was visualised in the interstitial space under the skin of the mice.

### Matrigel Plugs

Matrigel plugs with cells were prepared on ice by re-suspending 2×10^6^ healthy, HA or LV-FVIII HA BOECs in 500 µl of Matrigel (Corning). Matrigel plugs without cells were prepared on ice by using 500 µl of Matrigel (Corning). The mixture was implanted intradermally in 8-week-old NSG or NSG- HA mice. For FVIII stimulation, 3 IU/ml of rhFVIII was added to Matrigel and 2 IU of rhFVIII were injected within the Matrigel plug every 2 days. After 10 days, the plugs were removed, fixed, and embedded in paraffin for histological analysis.

### Statistical Analysis

All data were expressed as mean ± SD. Graphs were generated, and statistical analyses were performed using Prism 8 (Graph Pad). The p values were calculated using the Student’s t test with a two-tailed distribution, assuming an equal standard deviation. Where described, a one-way ANOVA were performed with a Bonferroni post-hoc test; p < 0.05 values were considered statistically significant. ∗p < 0.05, ∗∗p < 0.01, ∗∗∗p < 0.001, ∗∗∗∗p < 0.0001.

## Supporting information

Supplemental Results and Material and Methods

## Acknowledgments

The authors thank Roberta Annamaria Cirsmaru for lentiviral vector production and Prof. Angelo Lombardo for help to design the gRNA for CRISPR/Cas 9 system.

## Source of Founding

AF was supported by Telethon grant no. GGP19201, in part by Vanguard grant no. 874700 and by Ministero della Sanità GR-2018-12366399.

SO was supported by Telethon grant no. GGP19201 and by AIRC IG 2022 ID 27155.

CO was supported by Bando per la Ricerca Roche, Ministero della Sanità GR-2018-12366399

CB was supported by by Fondazione Cariplo grant No. 2018-0253

## Authors Contribution

CO, AC and IM designed and performed experiments and analysed data. SA, CS, GW and SM performed experiments and analysed data. CB conducted the in vivo studies. IM, FA, AL designed and performed RNA-Seq transcriptomic analysis. PC generated tools for CRISPR/Cas9 experiments. AF conceived the study. AF and SO generated funding analysed data and supervised the whole project. CO, AC, AF and SO wrote the paper that was revised by all authors and approved the final version.

