## Supplemental Results and Material and Methods for "FVIII interacts with cell surface to regulate endothelial cell functionality"

**Olgasi et al.**

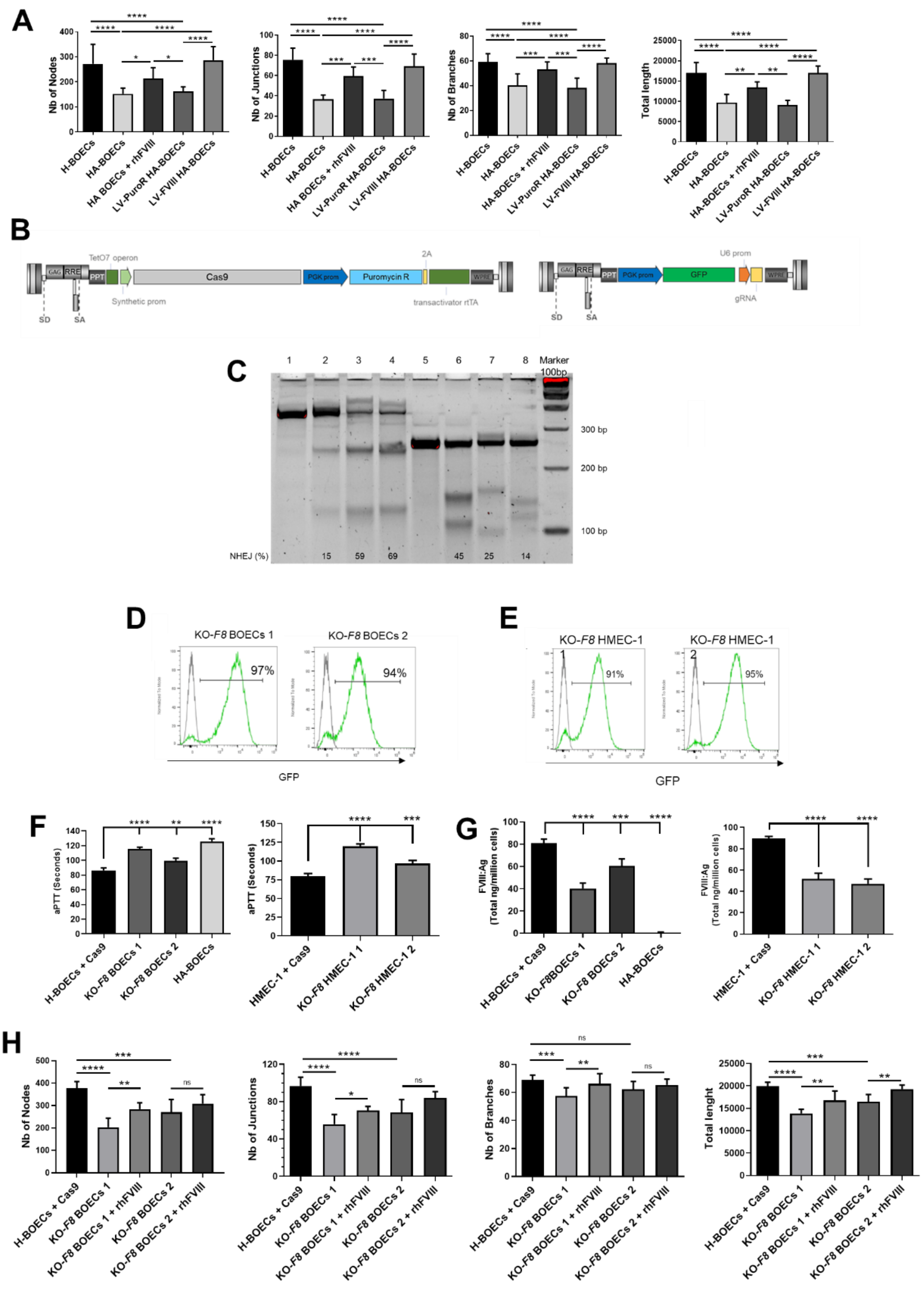


**Figure S1. HA-BOECs and KO-*F8* ECs are defective in tubulogenesis and migration. A)** Quantification of number of nodes, junctions, branches and total length of the tubule networks of H-BOECs (n=4), HA-BOECs (n=4) in presence or absence of rhFVIII, and LV-FVIII HA-BOECs (n=4). LV-PuroR was used as control for LV transduction. (****p<0.0001; ***p<0.001; **p<0.01; *p<0.05). Data are expressed as mean ± SD and are representative of three independent experiments. **B)** H-BOECs and HMEC-1 cells were transduced with a third generation LV carrying Cas9 under the control of a doxycycline inducible synthetic promoter and the puromycin resistance gene under the control of PGK promoter. A second LV construct, carrying the gRNA sequence under the control of U6 promoter and the GFP under the control of PGK promoter, was used to transduce ECs + Cas9 **C)** T7 endonuclease I cleavage assay performed on HEK293T cells transfected with 3 gRNA against exon 4 (lanes 2-4) or exon 7 (lanes 6-8) of *F8* to identify the best gRNAs that produce higher NHEJ. **D**) Representative histograms for GFP evaluation by FACS analysis in KO-*F8* BOECs 1 and 2 and **E)** KO-*F8* HMEC-1 1 and 2. **F)** aPTT assay on supernatant of KO-*F8* BOECs and KO-*F8* HMEC-1. HA-BOECs and ECs + Cas9 were used as controls. **G)** FVIII antigen assay on supernatant of KO-*F8* BOECs and KO-*F8* HMEC-1. HA-BOECs and ECs + Cas9 were used as controls. **H)** Quantification of number of nodes, junctions, branches and total length of the tubule networks of H-BOECs + Cas9, KO-*F8* BOECs 1 and 2 in presence or absence of rhFVIII. (****p<0.0001; ***p<0.001; **p<0.01; *p<0.05). Data are expressed as mean ± SD and are representative of three independent experiments.

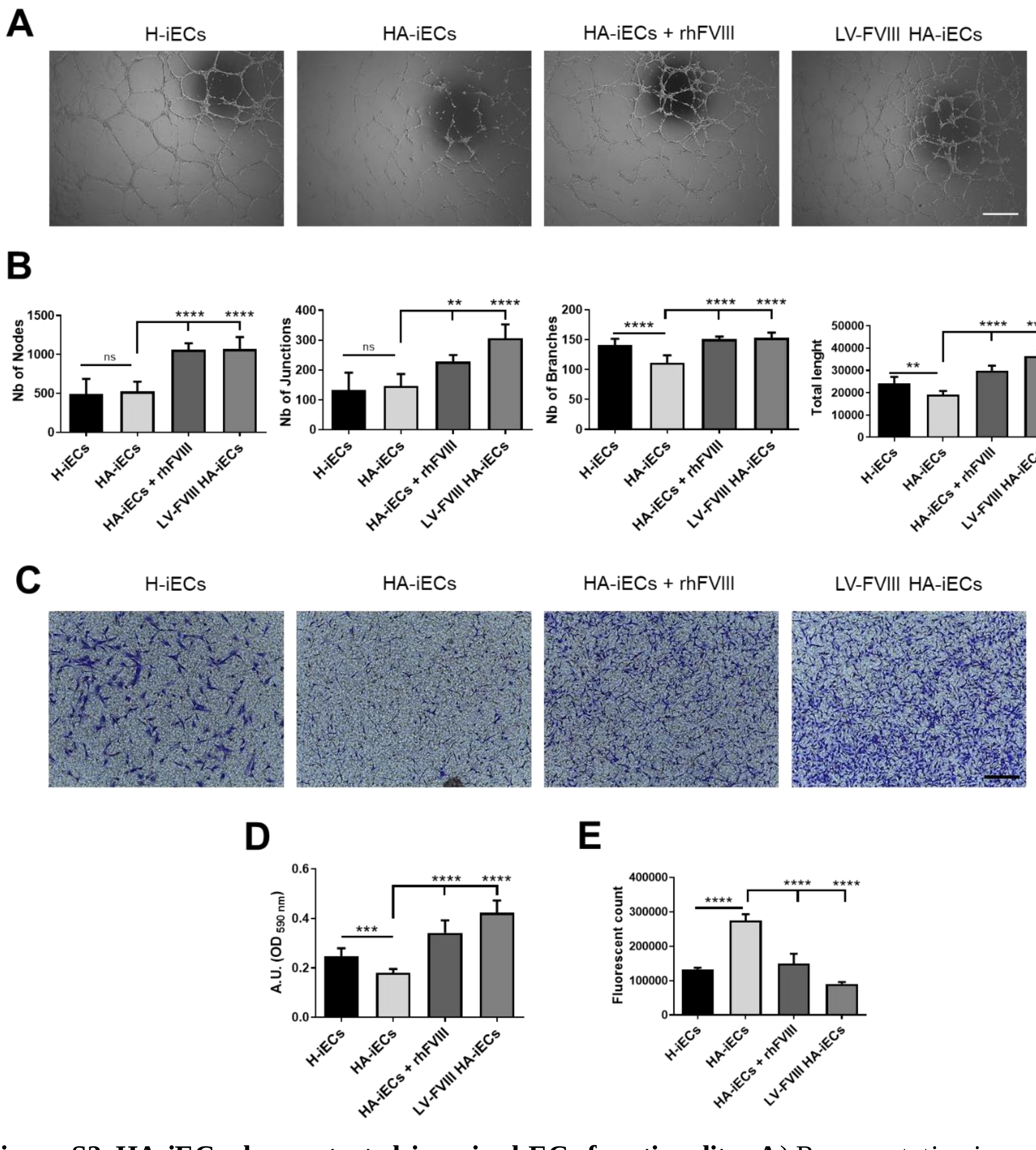


**Figure S2. HA-iECs demonstrated impaired ECs functionality. A**) Representative images of tubulogenic assay on H-iECs (n=2), HA-iECs (n=2) treated or not with rhFVIII and LV-FVIII HA-iECs (n=2). Scale bar = 500 µm. **B)** Quantification of number of nodes, junctions, branches and total length of the tubule networks; **C)** Representative images of migrated cells on H-iECs, HA-iECs treated or not with rhFVIII and LV-FVIII HA-iECs. Scale bar = 200 µm. **D)** Indirect quantification of cell migration assay by elution of crystal violet staining. **E)** Quantification of the extravasation of FITC-dextran through an intact monolayer on H-iECs, HA-iECs treated or not with rhFVIII and LV-FVIII HA-iECs (****p<0.0001; ***p<0.001; **p<0.01). Data are expressed as mean ± SD and are representative of three independent experiments.

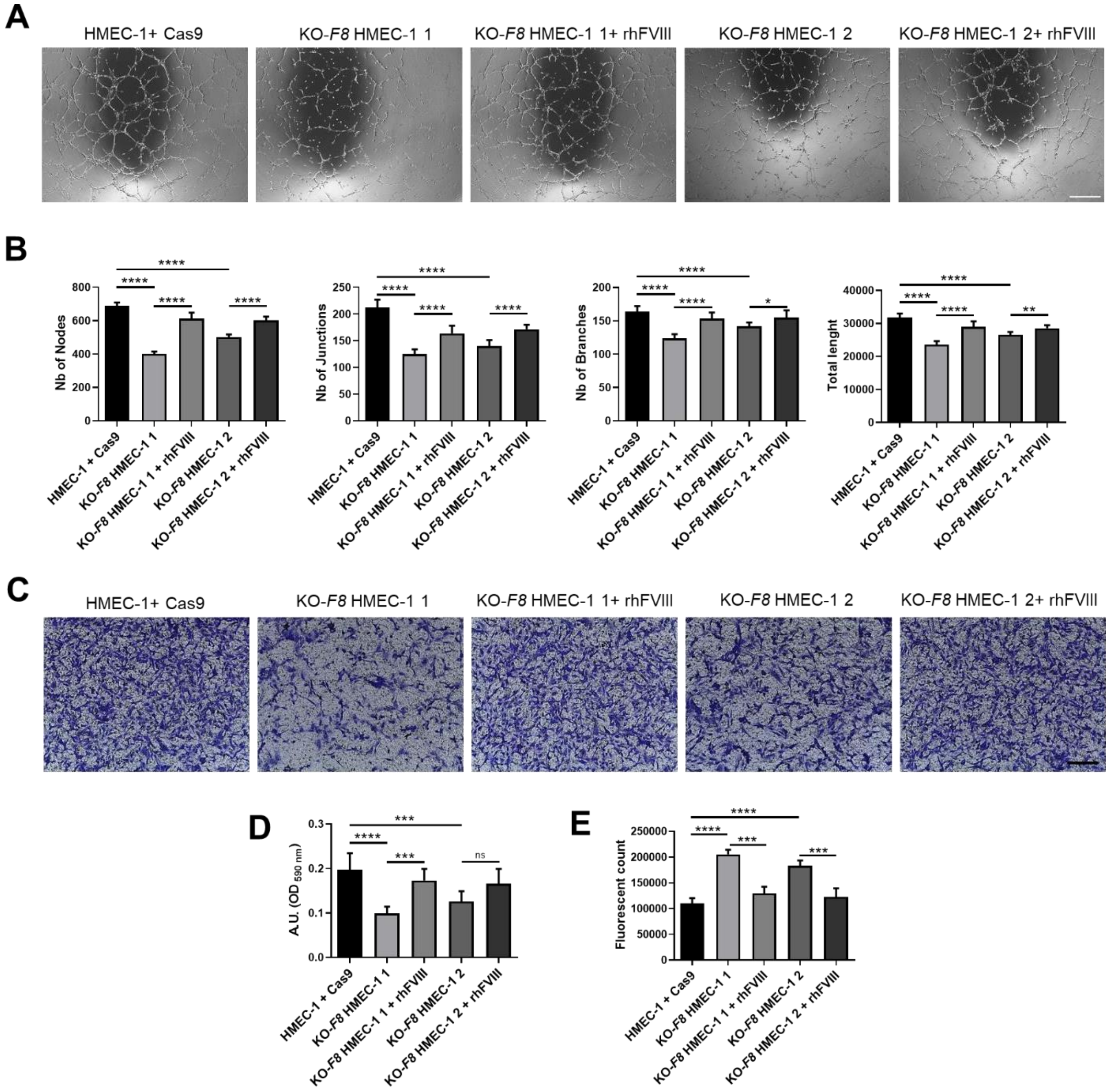


**Figure S3. FVIII knockout on HMEC-1 induces ECs instability. A**) Representative images of tubulogenic assay on HMEC-1 + Cas9, KO-*F8* HMEC-1 1 or 2 treated or not with rhFVIII. Scale bar = 500 µm **B**) Quantification of number of nodes, junctions, branches and total length of the tubule networks; **C**) Representative images of migrated cells on HMEC-1 + Cas9, KO-*F8* HMEC-1 1 or 2 treated or not with rhFVIII. Scale bar = 200 µm. **D**) Indirect quantification of cell migration assay by elution of crystal violet staining. **E**) Quantification of the extravasation of FITC-dextran through an intact monolayer on KO-*F8* HMEC-1 1 or 2 treated or not with rhFVIII. (****$p < 0.0001$; ***$p < 0.001$; ** p 0.01; * $p < 0.05$). Data are expressed as mean ± SD and are representative of three independent experiments.

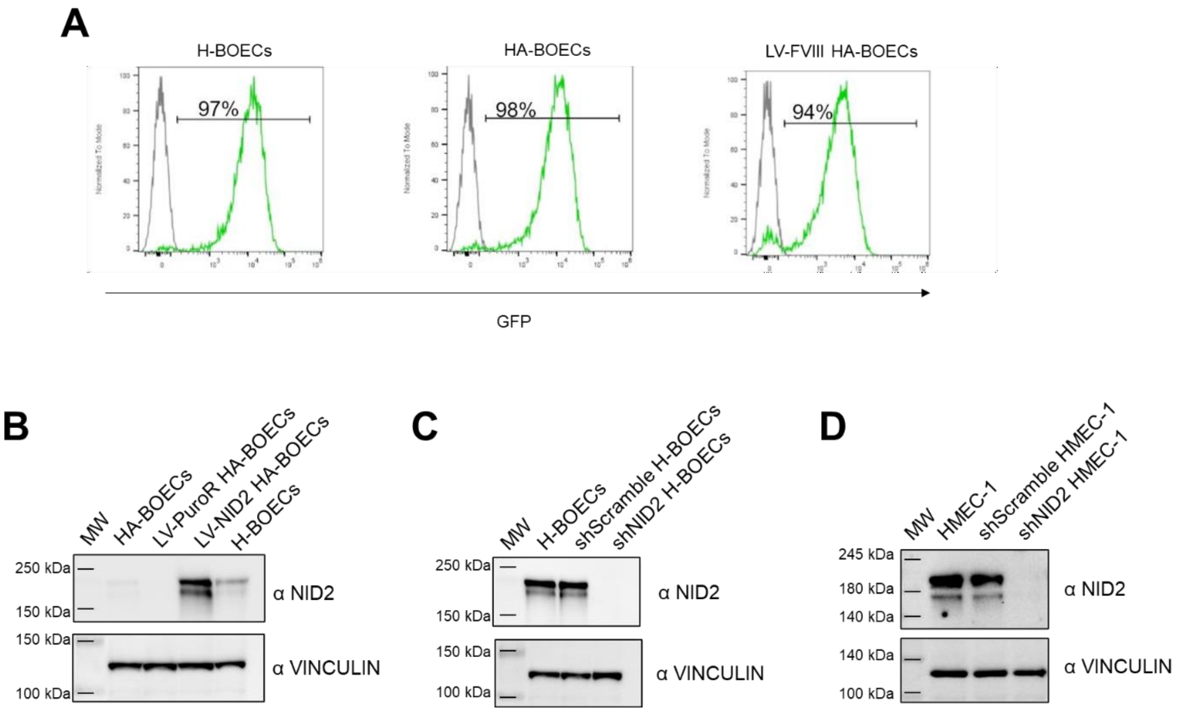


**Figure S4. LV transduction of BOECs and HMEC-1. A**) Representative FACS histograms showing GFP expression of H-BOECs, HA-BOECs and LV-FVIII HA-BOECs transduced with LV-GFP. The gray line of the histograms represents non-transduced BOECs. **B**) Western blot analysis of NID2 protein level on LV-NID2 HA-BOECs compared to HA-BOECs. LV.PuroR HA-BOEcs and H-BOECs were used as control. **C**) Western blot analysis of NID2 expression on shNID2 H-BOECs compared to non-transduced and shScramble H-BOECs. **D**) Western blot analysis of NID2 expression on shNID2 HMEC-1 compared to non-transduced and shScramble HMEC-1. In **B**, **C** and **D** Vinculin was used as loading control.

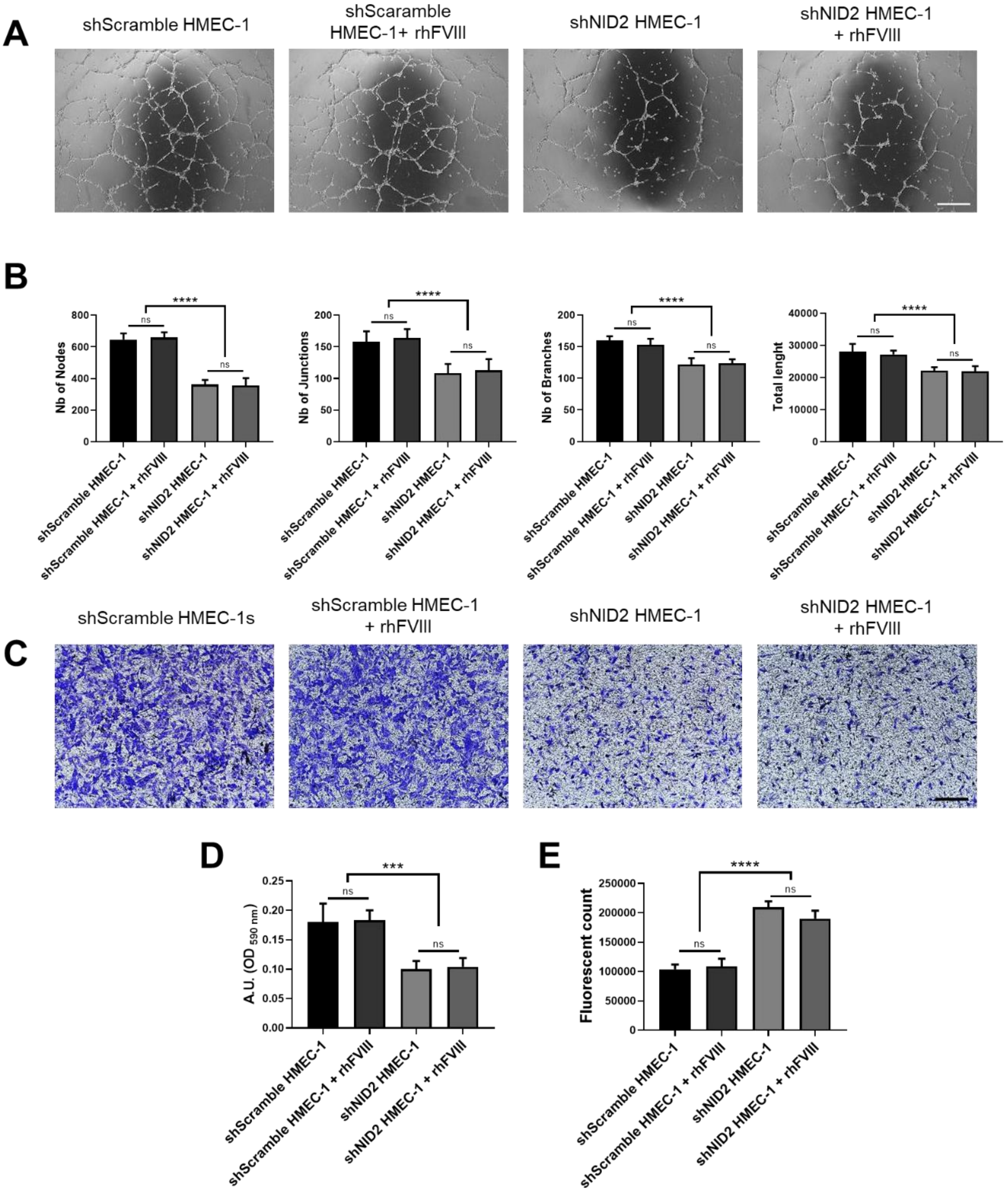


**Figure S5. NID2 knockdown impairs HMEC-1 endothelial functionality. A**) Representative images of tubulogenic assay on shScramble and shNID2 HMEC-1 cells in presence or absence of rhFVIII. Scale bar = 500 µm. **B**) Quantification of number of nodes, junctions, branches and total length of the tubule networks. **C**) Representative images of migrated cells on shScramble and shNID2 HMEC-1 cells in presence or absence of rhFVIII. Scale bar = 200 µm. **D**) Indirect quantification of migrated cells by elution of crystal violet staining. **E**) Permeability assay quantification calculated on the extravasation of FITC-dextran through an intact monolayer of shScramble and shNID2 HMEC-1 cells in presence or absence of rhFVIII. (****p < 0.0001***p < 0.001). Data are expressed as mean ± SD and are representative of three independent experiments.

### Supplemental Methods

#### Cell culture

HEK293T cells were cultured in Iscove's Modified Dulbecco's medium (IMDM) (Gibco) supplemented with 10% fetal bovine serum (Gibco) and 1% Penicillin-Streptomycin (100U/ml Euroclone) under humidified atmosphere of 5% CO2 at 37 °C.

HMEC-1 (human dermal microvascular endothelial cell line) (ATCC, CRL-3243) were cultured in MCDB131 medium (Gibco) supplemented with 10% fetal bovine serum (Gibco), 10 ng/ml human Epidermal Growth Factor (Immunotools), 1 μg/ml hydrocortisone, 1% Glutamine (Euroclone) and 1% Penicillin-Streptomycin (100U/ml Euroclone).

#### Lentiviral vector generation.

Third generation self-inactivating LVs were produced as previously published (1). Briefly, HEK293T cells were expanded and transiently transfected by the calcium phosphate precipitation method with four plasmids encoding for two core packaging constructs (pMDLg/pol and pRSV-Rev), the envelope construct (pMD.VSV.G), and the transfer vector construct (pVEC.FVIII; pPGK.FVIII; pPGK.PuroR; pPGK.NID2; pPGK.shScramble; pPGK.shNID2; pF8P.FVIII; pVEC.GFP; pPGK.Cas9; pPGK.GFP.U6promoter.gRNAex4F8; pPGK.GFP.U6promoter.gRNAex7F8). The cell supernatant was harvested, and LV particles were concentrated by ultracentrifugation. The GFP expressing LVs were titrated by flow cytometry analysis (FACS), while the other LVs were quantified as number of integrated copies by quantitative-PCR (qPCR) as previously described (2).

#### Cell transduction

HA-BOECs were plated at a $10^4$ cells/cm$^2$ density and after 6–8 h transduced with a lentiviral vector carrying the BDD form of FVIII.2A.PuroR under the control of the ubiquitous PGK promoter, or a LV carrying the Nidogen2.2A.PuroR or Puromicin Resistance gene under the control of PGK promoter using a Multiplicity of infection (MOI) of 20. H-BOECs, HA-BOECs and LV-FVIII BOECs were LV transduced with a LV carrying the GFP under the control of the VE-cadherin promoter (LV-VEC.GFP), using an MOI of 20 for Matrigel plugs experiments. HA-ECs were transduced as previously described (3). H-BOECs and HMEC-1 were transduced with LV.Cas9 and later with LV.PGK-GFP.U6promoter.gRNA with an MOI of 10. HMEC-1 were LV transduced also with LV.shNID2 or LV.shScramble with a MOI of 10. After 14–16 h incubation, fresh medium was added to the cells and, 72 h later, half of the cells were harvested for subsequent analysis, while the other half was further cultured.

**Guide RNA design and cloning**

gRNAs against the human F8 gene were designed using the website tool CHOP CHOP (https://chopchop.cbu.uib.no). Three guides showing high activity score and no predicted off-target activity were selected for both exon 4 and 7 (Table S1). The CHOP CHOP tool was used to design primers to amplify the regions encompassing the Cas9-cleavage sites. The gRNA sequences (Table S2) were ordered as oligo by Sigma Aldrich, with appropriate overhangs, annealed and cloned by BbsI digestion (New England BioLabs) into a LV transfer construct containing the U6 gRNA-expression cassette into the self-inactivating Long Terminal Repeats (LTRs) and encoding for the marker gene GFP from the human PGK promoter (pCCLsin.cPPT.hPGK.eGFP.Wpre.3'LTR-U6.gRNA loxP or LV-Cas9). Concerning the cloning of the Cas9-expressing LV, it was generated by replacing the Cas9 promoter of the pCW-Cas9 plasmid (Addgene No. 50661) with the SK-T6 promoter (4).

**gRNA selection in HEK293T cells**

The LV plasmids coding for the gRNAs were individually transfected with a Cas9-expressing plasmid (5) in HEK293T cells by electroporation using the 4D-Nucleofector™ System (Lonza, Basel, Switzerland) following the manufacturer's instructions. Briefly, HEK293T were cultured to 60–70% confluence, then harvested and washed with phosphate buffered saline. Approximately $5\times10^5$ cells were resuspended in the nucleofection solution SF together with 1µg of the Cas9 plasmid and 500 ng of the gRNA-expressing plasmid and transfected using instrument's program CM-130. 10 days post-nucleofection, genomic DNA from transfected cells was extracted using Maxwell 16 LEV Blood DNA kit (Promega) and the region encompassing the Cas9 cleavage sites in exon 4 or 7 were PCR amplified using primes listed in Table 3 and according to this conditions: 30 cycles of heating at 95 °C for 30 s, 58 °C for 30s, and 72°C for 1 min, with a final extension of 2 min at 72°C amplify. PCR amplicons were then subjected to the T7 endonuclease mismatch assay (New England Biolabs) according to manufacturer's instruction. The T7-treated PCR products were resolved using High Resolution Precast Electrophoresis Gel Spreadex® (AL-Diagnostic-GMBH) and visualized by staining with Atlas ClearSight Dna Stain (BioAtlas) using the Gel Doc System (Biorad). Cas9-induced mutations at exon 4 and 7 was quantified as previously described (5) (Figure. S1C). Based on these data, Ex4 gRNA 3 and Ex7 gRNA1, which resulted in the higher editing efficiency, were chosen for further experiments.

**Flow cytometry analysis**

Healthy, HA and LV-FVIII HA-BOECs transduced with LV-GFP and H-BOECs and HMEC-1 transduced with pPGK.GFP.U6promoter.gRNAex4F8 or pPGK.GFP.U6promoter.gRNAex7F8 were analyzed to assess GFP positivity. For each sample, $1.5\times10^5$ live events were acquired on the Attune NxT Acoustic Focusing Cytometer

(ThermoFisher Scientific, Waltham, MA, USA). Data were analyzed by FlowJo V10 software (BD Biosciences).

**FVIII Activity and FVIII Antigen Assay**

To assess FVIII activity, aPTT assay was performed on supernatants of cultured cells. Standard curves were generated by serial dilution of recombinant human FVIII (rhFVIII) in culture medium. Samples and standards were diluted 1:5 in FVIII-deficient plasma (TECO). For *in vivo* experiments, aPTT assay was performed on plasma samples of WT, HA and LV-injected mice and standard curves were generated by serial dilution of rhFVIII in hemophilic mouse plasma. All the experiments were performed using a Coatron M4 coagulometer (TECO Medical Instruments) and TEClot APTT-S kit reagents (TECO Medical Instruments).

FVIII antigen in supernatant of cultured cells was quantified by ELISA sandwich using the Matched-Pair Antibody Set for ELISA of human Factor VIII antigen (Affinity Biologicals) following the manufacturer's protocol. For standard curves generation, known concentrations of commercial rhFVIII were serially diluted in culture medium.

**Immunostaining**

Mouse Matrigel plugs were harvested and fixed in 4% PFA for 2h at 4°C and embedded in cryostat embedding medium (Bio-Optica). Cryostat sections of 4µm thickness were blocked in buffer containing 5% goat serum, 1% BSA, and 0.1% Triton X-100 in PBS, incubated with primary antibody followed by the secondary antibody at RT. For nuclei detection DAPI was added to the secondary antibodies' solution. Stained cells were analyzed using a Leica DM5500 microscope and images were acquired using the Leica application suite X software (LAS-X). Primary and secondary antibodies and dilutions are reported in Table 4.

**Western Blot**

Whole cell lysates were prepared using RIPA buffer (50 mmol/L Tris pH 7.5, 150 mmol/L NaCl, 1% NP40, 1% sodium deoxycholate, 0.1% SDS, 1× protease inhibitor cocktail (Sigma Aldrich)) with concentrations determined using the Pierce BCA Protein Assay (Thermo Fisher Scientific). The samples were size-fractionated on 8% SDS-PAGE under reducing conditions and electro-transferred to immuno-blot polyvinylidene difluoride membrane (BioRad). Membranes were incubated with anti-NID2 antibody (1:3000, Invitrogen) and anti-Vinculin antibody (1:10000, SantaCruz) and visualized with the appropriate horseradish peroxidase-conjugated secondary antibody. Immunoreactive proteins were detected using enhanced chemiluminescence (Clarity Western ECL Substrate; Bio Rad) with image capture performed using ChemiDoc Touch Imaging System (BioRad).

**Animal procedures**

Animal studies were approved by the Animal Care and Use Committee at UPO (Italian Health Ministry Authorization nos. 492/2016-PR and DBO64.5). NOD.Cg-Prkdc$^{scid}$Il2rg$^{tm1Wjl}$/SzJ (Jackson stock No 005557) mice were purchased by Charles River while the NSG-HA mice were previously generated and maintained in our laboratory (6). All animals' procedures were performed under a sterile hood.

**Table S1. Sequences of the crRNA present in the gRNAs**

| | |
|---|---|
| Ex4 gRNA 1 | GTACTAGTAGGGCTCCAATG |
| Ex4 gRNA 2 | GGACCTGCCAGACATATGTA |
| Ex4 gRNA 3 | GTGGAAGCCATACATATGTC |
| Ex7 gRNA 1 | GCTTGTGAGGAACCATCGCC |
| Ex7 gRNA 2 | GAACCATCGCCAGGCGTCCT |
| Ex7 gRNA 3 | GTGCACTCAATATTCCTCGA |

**Table S2. Oligos for gRNA cloning**

| | |
|---|---|
| F8.Ex4_1f | ACCGTACTAGTAGGGCTCCAATG |
| F8.Ex4_1r | AAACCATTGGAGCCCTACTAGTA |
| F8.Ex4_2f | ACCGGACCTGCCAGACATATGTA |
| F8.Ex4_2r | AAACTACATATGTCTGGCAGGTC |
| F8.Ex4_3f | ACCGTGGAAGCCATACATATGTC |
| F8.Ex4_3r | AAACGACATATGTATGGCTTCCA |
| F8.Ex7_1f | ACCGCTTGTGAGGAACCATCGCC |
| F8.Ex7_1r | AAACGGCGATGGTTCCTCACAAG |
| F8.Ex7_2f | ACCGAACCATCGCCAGGCGTCCT |
| F8.Ex7_2r | AAACAGGACGCCTGGCGATGGTT |
| F8.Ex7_3f | ACCGTGCACTCAATATTCCTCGA |
| F8.Ex7_3r | AAACTCGAGGAATATTGAGTGCA |

**Table S3. Oligo for the amplification of exon 4 and 7 of F8**

| | |
|---|---|
| F8.Ex4.T7_F | TTGAGTGTACAGTGGATATAGAAAGG |
| F8.Ex4.T7_R | TCAGGTGAAGGAACACAAATGC |
| F8.Ex7.T7_F | TCATAGCCATAGGTGTCTTATTCC |
| F8.Ex7.T7_R | ATGTTGGTGGGAAGAGATATGAC |

**Table S4. Primary and secondary antibodies**

| Antigen | Reactivity | Manufacturer | Format |
|---|---|---|---|
| CD31 | mouse | Clone Mec 13.3 Biolegend | Biotin |
| αSMA | mouse | Clone 1A4 Abcam | Not conjugated |
| / | / | Streptavidin eBioscience | PE |
| GFP | / | Polyclonal Thermo Scientific | Not conjugated |
| NID2 | human | Polyclonal Thermo Fisher | Not conjugated |
| Vinculin | human | Clone H-10 Santa Cruz | Not conjugated |
| IgG (H + L) | mouse | Polyclonal Thermo Scientific | 488 |
| IgG (H + L) | mouse | Polyclonal Thermo Fisher | HRP |
| IgG (H + L) | rabbit | Polyclonal Thermo Fisher | HRP |
